# Selective State Space Models Outperform Transformers at Predicting RNA-Seq Read Coverage

**DOI:** 10.1101/2025.02.13.638190

**Authors:** Ian Holmes, Johannes Linder, David Kelley

**Affiliations:** Department of Bioengineering, University of California, University Drive, Berkeley 94703; Calico Life Sciences LLC, 1170 Veterans Blvd, South San Francisco, CA 94080

## Abstract

Transformers are the basis for many state-of-the-art machine learning tools, including those for predicting gene expression data from DNA sequence. The considerable time and cost of training transformer models has motivated development of alternative approaches inspired by ideas from the signal-processing literature, such as state-space models (Mamba), Fourier transforms (Hyena), and wavelet transforms (MultiResNet). To evaluate these methods as potential replacements (or complements) for attention, we developed a software library bilby, implemented using Python and Jax/Flax, providing convolutional, attention, bidirectional Hyena, bidirectional Mamba, and striped-architecture models for supervised multi-task learning in functional genomics. We report a comparison of these architectures, testing several hyperparameters and variations, and reporting performance statistics for the withheld test set as well as downstream SNP classifiers. Relative to models comprising convolution and attention layers (implemented in Python and TensorFlow via the Baskerville library used by the Borzoi software), models comprising convolutional, bidirectional Mamba, and (optionally) attention layers achieve small but consistent improvements in prediction accuracy, for roughly comparable training times and parameter counts, when averaged across all output tracks and data splits (a proportional increase of 3-4% in Pearson R, and 1-2% in r^2^, with the highest gains achieved when Mamba and attention layers were combined in a striped architecture). In contrast, Hyena (when reimplemented as described in the literature) was not competitive with attention-based models at these tasks, while MultiResNet proved too slow to be practical. The gains in prediction accuracy of the Mamba-based models do not yet translate to significantly improved performance on downstream SNP classification tasks: benchmarks using a GTEx eQTL dataset yield roughly similar results for Mamba- and attention-based classifiers, with attention marginally outperforming Mamba in one metric (a difference of +0.007 in area under ROC) and slightly underperforming by another metric (a difference of −0.006 in Spearman rank correlation). We argue that these results suggest selective state-space models (such as Mamba and Striped Mamba) warrant further exploration for functional genomics tasks. Our code and trained models are publicly available at https://github.com/ihh/bilby.

## Introduction

An ongoing challenge of functional genomics is to predict the effect of regulatory DNA sequences on the control of gene expression. Success at this task has many downstream applications including, for example, classification of the deleterious impact of variants (Linder et al. 2025), identification of binding sites for regulatory proteins (Shrikumar et al. 2018), and design of regulatory sequences with desired properties (Schreiber, Lu, and Noble 2020).

In machine learning terms, this is a seq2seq problem where the input sequence is the genomic DNA and the output a stacked set of genome browser tracks, representing experimental data from large-scale functional genomics collections such as ENCODE. To ensure fixed-size output tensors, these aligned reads are typically summarized by their sequence coverage and aggregated into fixed-size bins at some resolution, yielding a single vector for each output track. Performance is improved by incorporating multiple types of functional genomics experiment in the output such as RNA-Seq, CAGE, DNase-Seq, ATAC-Seq, and ChIP-Seq (Kelley 2020; Kelley et al. 2018; Avsec et al. 2021; Linder et al. 2025), which highlight a wide set of salient gene and chromatin factors.

The earliest neural models for this task used CNNs (Kelley et al. 2018), with later models such as Enformer adding attention layers to better integrate long-range interactions (Avsec et al. 2021). The current state-of-the-art method Borzoi uses a similar architecture to Enformer including both convolution and attention layers (Linder et al. 2025). The convolution layers act at the earliest stages to detect local motifs, e.g. binding sites for transcription factors or other proteins, and local clusters of such motifs. The attention layers operate at later stages and model the coordinated effect of combinations of such regions over longer scales. Attention significantly boosts predictive accuracy, but it is a resource-intensive mechanism: every position in a length-L sequence must attend to every other position, requiring O(L^2^) operations. A naive implementation of attention thus requires O(L^2^) time and memory, and this limits the context length that can be used in such models. In order to mitigate this, the Borzoi attention layers operate at a lower resolution (128bp) than the final output resolution (32bp); the output of the attention layers is up-sampled back to 32bp resolution using the U-Net architecture (Ronneberger, Fischer, and Brox 2015).

Several techniques to make attention more efficient have been proposed. The O(L^2^) operations can be performed in parallel (Liu, Zaharia, and Abbeel 2023), or can be scheduled in such a way as to take optimal advantage of GPU memory caches and bus structure (Dao et al. 2022). However, these techniques mitigate the O(L^2^) problem, rather than eliminating it entirely, and come with drawbacks: parallelization demands more GPUs, while cache management entails tricky low-level CUDA programming. Other approaches, such as low-rank approximations (S. Wang et al. 2020), random basis projections (Choromanski et al. 2020), or locality-sensitive hashing (Kitaev, Kaiser, and Levskaya 2020) reduce time complexity but at a cost in accuracy and/or predictability. Consequently, training deep transformers remains time-consuming and expensive, which raises the barrier to new model development and potentially impedes downstream fine-tuning.

The training cost of transformers has motivated efforts in the ML community to explore sub-quadratic alternatives to attention, including dilated convolutions such as WaveNet (van den Oord et al. 2016) and MultiResNet (Shi, Wang, and Fox 2023), state space models (SSMs) such as H3 (Fu et al. 2022), the Selective SSM Mamba (Gu and Dao 2023), and the SSM-adjacent Hyena (Poli et al. 2023). Many of these are based on techniques from signal processing and function approximation: MultiResNet is inspired by wavelet transforms, Hyena by Fourier transforms, and SSMs by a family of methods including linear differential equations for physical systems (such as those used in electrical circuit analysis or Newtonian dynamics), autoregressive moving-average models, and orthogonal polynomial basis approximations.

The SSM-based approaches have generated especially promising results, especially when SSM layers are alternated with attention layers in what has been called a “striped” architecture (Nguyen et al. 2023; Schiff et al. 2024). Striped SSM architectures have been applied successfully to several problems in genomics including unsupervised language modeling (Nguyen et al. 2023, 2024; Schiff et al. 2024) and to the generation of spliced reads (Fradkin et al. 2024).

Here, we report an investigation into the utility of MultiResNet, Hyena, Mamba, and striped Mamba as architectures for predicting RNA-seq coverage from DNA sequence data, the task performed by the Borzoi model (Linder et al. 2025). We assessed performance both directly on the modeling task itself, quantified by the correlation of predicted output with test data, and indirectly by passing the model’s outputs to a downstream SNP classifier. Our code is implemented using Jax, the differentiable Python framework, and includes fresh or newly-modified implementations of several of the novel methods (standard versions of these models were unavailable when we began this work; there are now more implementations available in the PyTorch library). The code is publicly available at [URL] along with our trained model parameters.

## Results

The models investigated in this study are listed in Table 1, together with their parameter counts and illustrative training times. Training curves for several of the models are shown in Figure 1, which plots the value of Pearson’s correlation coefficient (r), averaged across outputs on the validation set, during the progress of a training run for several of the models tested. As a comparison baseline, we have shown the performance for the convolutional trunk of the model, without any additional attention or other layers.

**Table 1.** Model architectures evaluated in this study include convolution (Kelley et al. 2018), attention (Vaswani et al. 2017; Avsec et al. 2021; Linder et al. 2025), MultiResNet (Shi, Wang, and Fox 2023), Hyena (Poli et al. 2023), Mamba (Gu and Dao 2023), and Striped Mamba (Schiff et al. 2024). All evaluated architectures included five initial rounds of convolution and twofold max-pooling leading to a 768-dimensional embedding over bins of 32-base resolution, i.e. a tensor of shape (12268,768); these layers are omitted from the descriptions shown in this table (see Figure 3 for examples). For the Mamba models, N is the hidden dimension and E is the expansion factor for the dense projection into the internal dimension of the Selective State Space Model (Gu and Dao 2023). Total training times are the time from commencement of training to the time of finding the best parameterization, averaged over all cases in which training reached a stopping point defined by validation set metrics. The exception is those cases for which training was stopped manually before this point was reached, in which case this is reported in the final column, and the reported training time is the average time at which the manual stop occurred. The final column indicates whether training completed (by reaching the validation set-defined stopping point), vs being stopped manually. All times are provided for illustrative comparison; in practice training times will depend on many software and hardware factors including implementation of backprop (vector-Jacobian product), GPU architecture, clock rate, performance of the Jax compiler or XLA engine, and random initialization.

| Model name | Codebase | Architecture | Number of parameters | Training time |  |  |
| --- | --- | --- | --- | --- | --- | --- |
|  |  |  |  | Hours/epoch | Hours total | Stopped? |
| CNN | bilby | Convolution only | 7,038,354 | 0.74 | 24 | No |
| Trans4 | Baskerville | [UNet + Transformer] x4 | 28,419,058 | 1.87 | 366 | Yes |
| MultiResNet | bilby | MultiResNet | 8,232,082 | 7.46 | n/a | Yes |
| Hyena4 | bilby | Hyena x4 | 16,640,274 | 1.70 | 102 | No |
| Hyena5 | bilby | Hyena x5 | 19,040,754 | 1.91 | 99 | No |
| Hyena6 | bilby | Hyena x6 | 21,441,234 | 2.19 | 105 | No |
| Hyena7 | bilby | Hyena x7 | 23,841,714 | 2.43 | 135 | Yes |
| Hyena8 | bilby | Hyena x8 | 26,242,194 | 2.68 | 146 | Yes |
| Mamba16 | bilby | Mamba(N=8,E=1) x6 | 22,374,546 | 3.87 | 352 | No |
| Mamba16-16 | bilby | Mamba(N=16,E=1) x6 | 22,595,922 | 6.19 | 223 | Yes |
| Mamba16-20 | bilby | Mamba(N=20,E=1) x6 | 22,706,610 | 8.05 | 353 | No |
| Mamba17 | bilby | Mamba(N=8,E=1) x7 | 24,930,578 | 4.38 | 346 | No |
| Mamba26 | bilby | Mamba(N=8,E=2) x6 | 37,696,146 | 7.21 | 347 | No |
| Mamba27 | bilby | Mamba(N=8,E=2) x7 | 42,805,778 | 7.35 | 347 | No |
| StripedMamba22 | bilby | [Mamba(N=8,E=1)x2 + UNet+Transformer+RoPE] x2 | 27,518,610 | 2.86 | 395 | No |
| StripedMamba22-20 | bilby | [Mamba(N=20,E=1)x2 + UNet+Transformer] x2 | 27,739,730 | 5.10 | 348 | No |
| StripedMamba32 | bilby | [Mamba(N=8,E=1)x3 + UNet+Transformer+RoPE] x2 | 32,630,674 | 3.70 | 305 | No |

**Figure 1.**
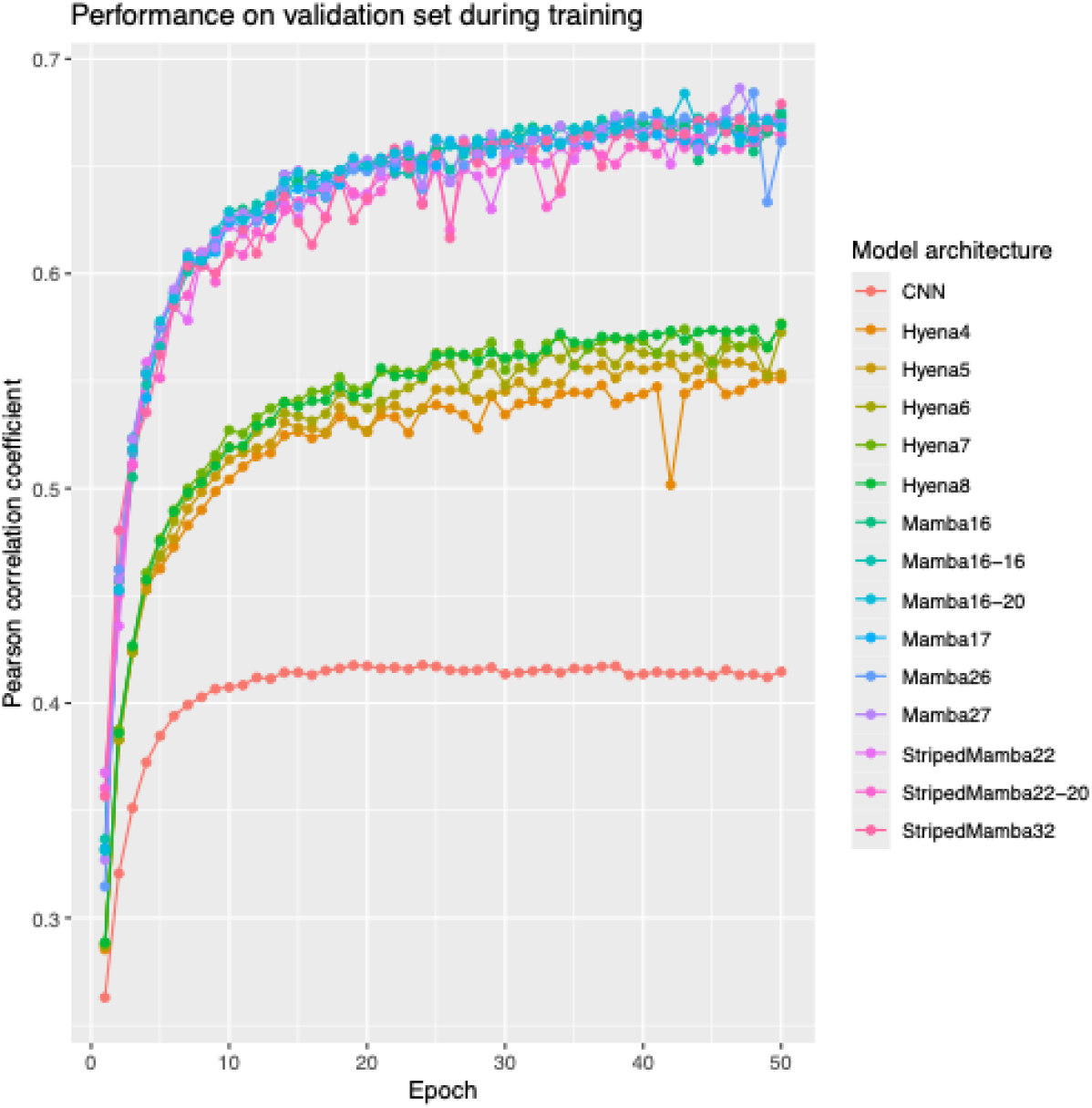
Progress through the first 50 epochs of training CNN, Hyena, and Mamba-based models as monitored by Pearson’s R on the validation data split reveals clear separation between convolution-only, Hyena-based, and Mamba-based models.

### MultiResNet

We began an evaluation of the wavelet-inspired MultiResNet, but abandoned this early on, since the model’s multiple dilated convolutional layers yielded extremely slow training times and apparently weak performance (judging by the early stages of training). While analyzing this model, we observed that, for single-channel convolutions (as used by MultiResNet and Hyena), a single long convolution can equivalently replace serial dilated convolutions (since the space of one-dimensional convolutions is closed under composition and dilation). After this observation, we focused our investigations of convolutional models on Hyena, which begins directly with a long convolution implemented efficiently in Fourier space and adds additional expressive power through data-dependent gating.

### Hyena

All transformer, Hyena, and Mamba layers in our models are bypassed by residual connections; thus, in theory, adding further layers should only cause performance to increase. In practice, at some depth, training stops working as effectively. In particular, we noted issues with training Hyena at depth. Some of these were traceable to the random initialization procedure used. In the original Hyena paper (Poli et al. 2023), the long convolution filter is initialized using a SIREN function, i.e. a multilayer network with sinusoidal activations that has been shown to be an effective general-purpose function approximator (Sitzmann et al. 2020); random initialization of this SIREN layer’s weights must be handled carefully when it is used as a long convolution, otherwise the variance of the output can explode, causing problems at depth.

After adopting this initialization scheme, we found that adding Hyena layers to this convolutional trunk did improve accuracy, with diminishing returns beyond around 8 layers. Since training time also increases with depth, we report subsequent results only for the 8-layer Hyena.

We explored several variations on the basic Hyena architecture, the most effective of which was to add channel-mixing to the innermost nearest-neighbor convolution. This did appear to improve performance; however, in view of the overall relatively weak performance of Hyena compared to Mamba, we did not investigate Hyena further. It is possible that more recent iterations of the Hyena model currently available in PyTorch would outperform the version that we evaluated.

### Mamba

The performance of the Mamba-based methods is in a class above the Hyena-based methods, as is clearly evident from the training curve (Figure 1). We found that Mamba performance increased as we added layers, up to 6 layers. Beyond this depth, peak predictive accuracy begins to drop off. This may be indicative of problems with the training schedule.

### Striped Mamba

In general, we observed that “striping” – alternating Mamba layers with transformer layers at some ratio – improved performance relative to using Mamba alone. In evaluating transformers on these data, we tried several kinds of relative positional encoding, including the geometrically-spaced threshold functions described by (Avsec et al. 2021) as well as the more recent rotary positional embeddings (Su et al. 2021). However, consistent with other work, we found that positional encodings did not significantly enhance the transformer’s performance when used in conjunction with Mamba layers – presumably because the state space models of Mamba are already capable, by their nature, of generating sinusoidal guide signals or injecting positional information in other ways.

### Equivariance

Several authors have described equivariant formulations for DNA-processing neural networks whose results, by construction, transform predictably when the input DNA sequence is reverse-complemented. These have been called “parameter-tied” (Shrikumar, Greenside, and Kundaje 2017; Zhou, Shrikumar, and Kundaje 2021) or “revcomp-equivariant” networks (Mallet and Vert 2024; Brown and Lunter 2019). We developed revcomp-equivariant versions of our convolutional stack, and our bidirectional Mamba is approximately equivariant by construction. However, in keeping with reports from prior authors (Zhou, Shrikumar, and Kundaje 2021; Brown and Lunter 2019), we did not find significant benefits to enforcing strict equivariance on the convolutional layers; in fact generally this led to a slight reduction in peak accuracy. We explored an alternate regularization approach to equivariance, weakly penalizing departures from equivariance in the loss, but this did not improve accuracy either. Tying the forward and reverse weight parameters of the dense projection used to map the input signal into Mamba’s internal space did significantly reduce the parameter count, so we continued to maintain this constraint, but we observed negligible effects on accuracy. Several recent papers from the Mamba developers report greater benefits of an equivariant formulation of Mamba in a bidirectional context (Schiff et al. 2024).

### Test set evaluations

Figure 2 shows performance of the best Mamba and Striped Mamba configurations on the validation dataset (in the first of our four splits), compared to the transformer, with each datapoint representing an output track. Performance varies significantly between the different types of experiment, with the CAGE output tracks generally predicted worse by Mamba (relative to the transformer) and dragging the Mamba average down; the striped version of Mamba appears to perform slightly better on these tracks.

**Figure 2.**
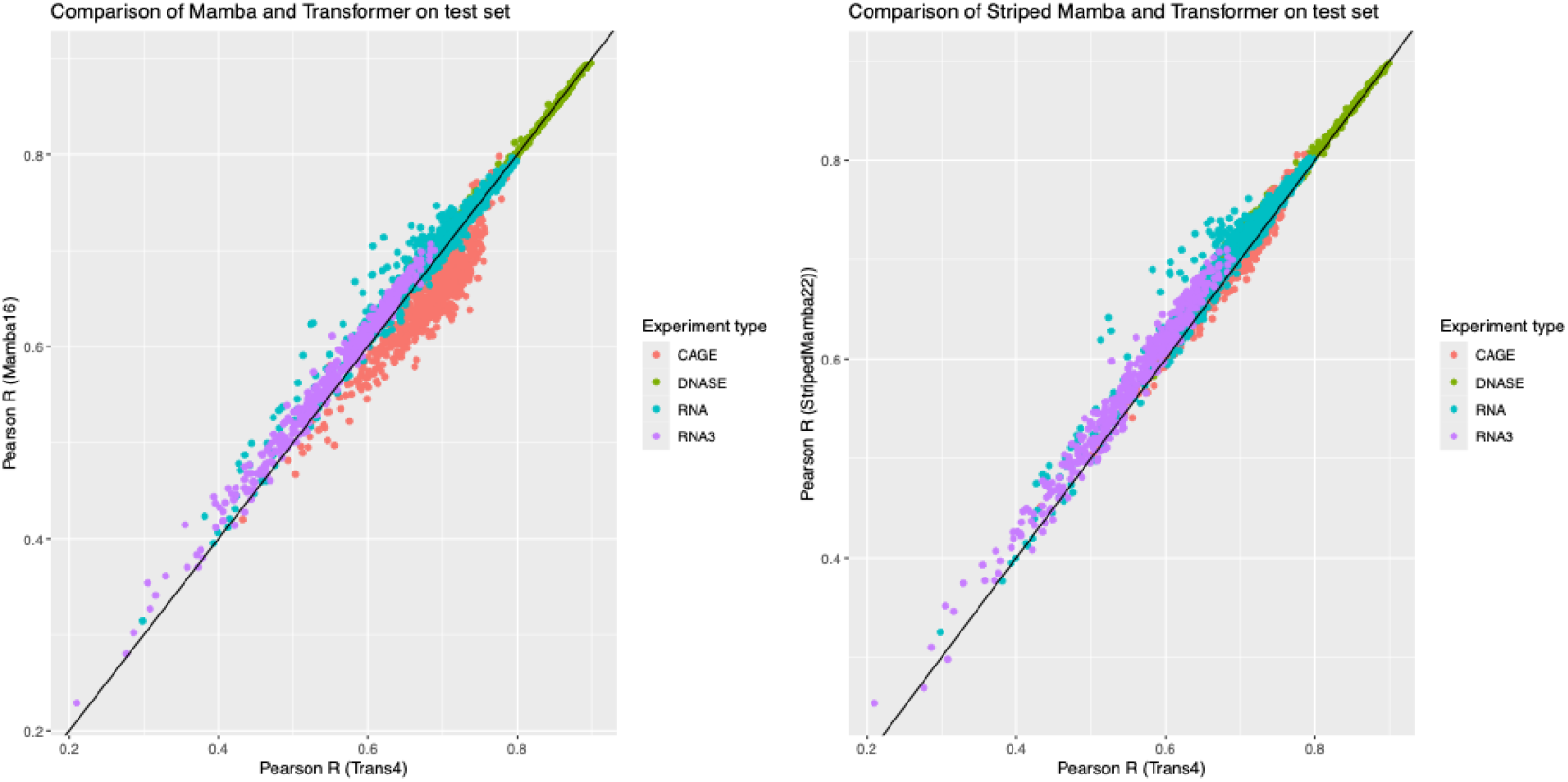
Comparison of Pearson correlation coefficients across the 2,194 output tracks for the 4-layer transformer-based model Trans4 (x-axis) with the 6-layer Mamba model Mamba16 (y-axis, left) and the striped Mamba model StripedMamba22 (y-axis, right) shows the close correspondence between the methods, with the Mamba model slightly outperforming the transformer. A breakdown by the experimental category of the output tracks shows that Mamba16’s worst performance is on the CAGE-Seq tracks, with most of the improvements by StripedMamba22 attributable to better performance on these tracks.

Table 2 shows the performance of the best methods on the test set, averaged across all four data splits. In general, Mamba achieves a small increase in prediction accuracy (as measured by Pearson R and r^2^) at comparable parameter counts, GPU usages, and total training times, an effect that is quite consistent and reproducible across different data splits and hyperparameter configurations (as long as the Mamba layers are stacked deep enough; our best models used 6 Mamba layers). We did not find that performance was improved by increasing the selective state space model’s hidden channel dimension (N) beyond 8. Nor was performance improved by increasing the “expansion factor” (E) projecting the model’s internal feature dimension into that of each Mamba layer; this factor is set at E=2 in the Mamba paper, but we found no improvement beyond E=1.

**Table 2.** Performance of models on the withheld test set show clear improvements for Mamba-based architectures over transformer architecture, for all types of output track. Performance is measured by Pearson R and coefficient of determination (R^2^), broken down by the output track’s experimental data type. All entries are averaged over four data splits. The final two columns show averages over tracks for all experiments.

| Model | DNase-Seq |  | CAGE-Seq |  | RNA-Seq |  | 3' RNA-Seq |  | All tracks |  |
| --- | --- | --- | --- | --- | --- | --- | --- | --- | --- | --- |
|  | R | R<sup>2</sup> | R | R<sup>2</sup> | R | R<sup>2</sup> | R | R<sup>2</sup> | R | R<sup>2</sup> |
| Trans4 | 0.7887 | 0.6220 | 0.6705 | 0.4398 | 0.6668 | 0.4453 | 0.5359 | 0.2829 | 0.6599 | 0.4373 |
| Hyena8 | 0.7529 | 0.5638 | 0.6283 | 0.3913 | 0.6107 | 0.3747 | 0.4456 | 0.2023 | 0.6043 | 0.3734 |
| Mamba16 | **0.7919** | 0.6271 | **0.6845** | 0.4641 | 0.6716 | 0.4503 | 0.5544 | 0.2994 | 0.6696 | 0.4500 |
| Mamba17 | 0.7900 | 0.6252 | 0.6876 | **0.4686** | 0.6708 | 0.4498 | 0.5535 | 0.3013 | 0.6691 | 0.4499 |
| Mamba26 | 0.7831 | 0.6099 | 0.6821 | 0.4616 | 0.6667 | 0.4444 | 0.5465 | 0.2906 | 0.6643 | 0.4428 |
| Mamba27 | 0.7804 | 0.6081 | 0.6774 | 0.4548 | 0.6663 | 0.4440 | 0.5491 | 0.2944 | 0.6632 | 0.4416 |
| StripedMamba22 | 0.7917 | **0.6279** | 0.6791 | 0.4575 | **0.6784** | **0.4634** | **0.5575** | 0.3122 | **0.6713** | **0.4557** |
| StripedMamba22-20 | 0.7802 | 0.6089 | 0.6736 | 0.4496 | 0.6723 | 0.4551 | 0.5529 | 0.3098 | 0.6649 | 0.4472 |
| StripedMamba32 | 0.7873 | 0.6222 | 0.6777 | 0.4532 | 0.6770 | 0.4559 | 0.5569 | **0.3142** | 0.6696 | 0.4527 |

This hyperparameter is the single most influential factor on the model’s parameter count (which scales roughly proportional to E).

### eQTL SNP classification

We also evaluated the model’s performance at a downstream SNP classification task, evaluating the various models’ abilities to distinguish fine-mapped GTEx expression quantitative trait loci (eQTLs) from a set of negatives. In this experiment, summaries of the model’s outputs were fed into a random forest classifier. Despite the reproducible gains in prediction accuracy obtained by using Mamba-based architectures, performance at the SNP classification task was not improved; in fact, we found (Table 3) that the transformer still slightly outperformed the Mamba-based methods at this task as measured by mean AUROC (0.7919 for the transformer, as compared to 0.7853 for the best Mamba-based method), although the Spearman rank correlation showed the opposite trend (0.2645 for the transformer, 0.2702 for the best Mamba-based methods). These numbers are extremely close; breaking the AUROC results down further, we find that the Mamba-based methods in fact outperform the transformer in 8 of the 49 tissues (Table 4). We also note that the mean AUROC for this small transformer-based model approaches that of the full Borzoi model (0.7943), although the Spearman rank coefficient reported for Borzoi is significantly better (0.3338). The closeness of these metrics is remarkable given that the model evaluated here is considerably smaller (Borzoi has an input window that is 524kb rather than 393kb, was trained on a larger dataset—7,611 human tracks and 2,608 mouse tracks rather than just 2,194 human tracks—using two 40Gb A100 GPUs rather than a 24Gb L4, uses an internal embedding dimension of 1,536 rather than 768, and contains 8 transformer blocks rather than 4).

**Table 3.** The transformer model outperforms other models on an GTEx eQTL SNP classification benchmark. Area under ROC curve (AUROC) and Spearman rank correlation of predictions are shown, using a random forest classifier that utilizes an ensemble of four models, one trained on each data split.

| Model | AUROC | Spearman |
| --- | --- | --- |
| Trans4 | **0.7919** | 0.2645 |
| Hyena8 | 0.7591 | 0.2030 |
| Mamba16 | 0.7853 | 0.2610 |
| Mamba17 | 0.7834 | 0.2316 |
| Mamba26 | 0.7826 | 0.2338 |
| Mamba27 | 0.7790 | 0.2356 |
| StripedMamba22 | 0.7838 | 0.2683 |
| StripedMamba22-20 | 0.7770 | 0.2692 |
| StripedMamba32 | 0.7812 | **0.2702** |

**Table 4.** In 41 out of 49 tissues, the transformer model outperforms other models on the GTEx eQTL SNP classification benchmark. Area under ROC curve (AUROC) and Spearman rank correlation of predictions are shown, using a random forest classifier trained on summary statistics of the outputs of models trained on all four data splits.

| Tissue | Trans4 | Hyena8 | Mamba16 | Mamba17 | Mamba26 | Mamba27 | StripedMamba22 | StripedMamba22-20 | StripedMamba32 |
| --- | --- | --- | --- | --- | --- | --- | --- | --- | --- |
| Kidney_Cortex | 0.759205 | 0.762913 | 0.783839 | 0.764527 | 0.784929 | 0.743769 | 0.782357 | 0.771978 | 0.772163 |
| Brain_Substantia_nigra | 0.783621 | 0.727337 | 0.749549 | 0.749434 | 0.759409 | 0.748444 | 0.745957 | 0.72511 | 0.74542 |
| Vagina | 0.799064 | 0.779303 | 0.799861 | 0.791506 | 0.808448 | 0.798198 | 0.805442 | 0.785364 | 0.800151 |
| Brain_Amygdala | 0.783024 | 0.743583 | 0.776669 | 0.768522 | 0.779375 | 0.750736 | 0.756449 | 0.765392 | 0.760228 |
| Uterus | 0.803482 | 0.741014 | 0.77602 | 0.771093 | 0.769557 | 0.775218 | 0.778781 | 0.757352 | 0.765743 |
| Brain_Spinal_cord_cervical_c-1 | 0.788675 | 0.746122 | 0.776137 | 0.769081 | 0.784491 | 0.76405 | 0.766936 | 0.763839 | 0.762134 |
| Brain_Anterior_cingulate_cortex_BA24 | 0.757246 | 0.725148 | 0.733882 | 0.733928 | 0.735159 | 0.733956 | 0.738581 | 0.734335 | 0.741592 |
| Minor_Salivary_Gland | 0.795485 | 0.760244 | 0.800608 | 0.787464 | 0.788192 | 0.775228 | 0.788982 | 0.769493 | 0.787029 |
| Cells_EBV-transformed_lymphocytes | 0.788405 | 0.741916 | 0.78132 | 0.778712 | 0.775642 | 0.772827 | 0.766507 | 0.76824 | 0.781862 |
| Brain_Hippocampus | 0.776125 | 0.740982 | 0.765938 | 0.767735 | 0.763893 | 0.754253 | 0.751039 | 0.761914 | 0.753768 |
| Brain_Hypothalamus | 0.813964 | 0.760443 | 0.790049 | 0.786404 | 0.78367 | 0.792137 | 0.796844 | 0.781436 | 0.78602 |
| Liver | 0.817712 | 0.76986 | 0.811171 | 0.807985 | 0.795715 | 0.796031 | 0.807937 | 0.798671 | 0.801364 |
| Ovary | 0.832839 | 0.778531 | 0.815935 | 0.826377 | 0.806622 | 0.81274 | 0.821902 | 0.807211 | 0.807699 |
| Small_Intestine_Terminal_Ileum | 0.813036 | 0.76867 | 0.792126 | 0.783134 | 0.785569 | 0.794423 | 0.808404 | 0.794972 | 0.801533 |
| Brain_Putamen_basal_ganglia | 0.793184 | 0.73333 | 0.782276 | 0.776429 | 0.775519 | 0.768123 | 0.774845 | 0.758027 | 0.77373 |
| Artery_Coronary | 0.80463 | 0.738887 | 0.789585 | 0.793839 | 0.779868 |  | 0.78382 | 0.770365 | 0.78342 |
| Brain_Frontal_Cortex_BA9 | 0.772211 | 0.733336 | 0.757544 | 0.758608 | 0.760001 | 0.755542 | 0.75202 | 0.752039 | 0.74849 |
| Prostate | 0.802986 | 0.762229 | 0.790448 | 0.789333 | 0.791573 | 0.779685 | 0.799021 | 0.781168 | 0.792091 |
| Brain_Nucleus_accumbens_basal_ganglia | 0.80976 | 0.758726 | 0.791548 | 0.795785 | 0.795881 | 0.793138 | 0.790657 | 0.788954 | 0.794566 |
| Brain_Caudate_basal_ganglia | 0.789959 | 0.735383 | 0.76021 | 0.76682 | 0.758736 | 0.760786 | 0.755826 | 0.762713 | 0.76131 |
| Adrenal_Gland | 0.814224 | 0.777064 | 0.810736 | 0.803802 | 0.804692 | 0.803397 | 0.802054 | 0.789763 | 0.800322 |
| Brain_Cortex | 0.809486 | 0.771732 | 0.789526 | 0.794803 | 0.792629 | 0.789054 | 0.78596 | 0.783824 | 0.781766 |
| Pituitary | 0.778156 | 0.739483 | 0.775975 | 0.760169 | 0.762236 | 0.765366 | 0.767753 | 0.769481 | 0.772339 |
| Brain_Cerebellar_Hemisphere | 0.809267 | 0.762232 | 0.79677 | 0.79782 | 0.794391 | 0.792025 | 0.794291 | 0.785333 | 0.791113 |
| Stomach | 0.807634 | 0.774944 | 0.801546 | 0.796838 | 0.797654 | 0.794204 | 0.800714 | 0.79577 | 0.804069 |
| Heart_Left_Ventricle | 0.792322 | 0.754366 | 0.785236 | 0.788558 | 0.781679 | 0.780897 | 0.783609 | 0.77872 | 0.775774 |
| Colon_Sigmoid | 0.79317 | 0.763605 | 0.796938 | 0.788608 | 0.795004 |  | 0.792425 | 0.778822 | 0.78754 |
| Brain_Cerebellum | 0.801812 | 0.762515 | 0.801497 | 0.801471 | 0.798386 | 0.791583 | 0.800206 | 0.792587 | 0.795439 |
| Pancreas | 0.806363 | 0.773513 | 0.795268 | 0.798364 | 0.789129 | 0.787332 | 0.795155 | 0.791119 | 0.794309 |
| Spleen | 0.818674 | 0.797301 | 0.8133 | 0.815041 | 0.813241 | 0.80904 | 0.817307 | 0.814187 | 0.81336 |
| Esophagus_Gastroesophageal_Junction | 0.797353 | 0.768733 | 0.787805 | 0.786233 | 0.781756 | 0.788844 | 0.786078 | 0.783289 | 0.784441 |
| Colon_Transverse | 0.791405 | 0.757473 | 0.783741 | 0.790875 | 0.787868 | 0.784586 | 0.785574 | 0.784998 | 0.785442 |
| Heart_Atrial_Appendage | 0.782991 | 0.757412 | 0.776338 | 0.777061 | 0.774565 | 0.769723 | 0.771443 | 0.767536 | 0.77434 |
| Breast_Mammary_Tissue | 0.803277 | 0.779235 | 0.79597 | 0.795173 | 0.790541 | 0.791292 | 0.797625 | 0.788721 | 0.789696 |
| Artery_Aorta | 0.798312 | 0.776322 | 0.788967 | 0.793316 | 0.787975 | 0.7929 | 0.79232 | 0.788824 | 0.786825 |
| Adipose_Visceral_Omentum | 0.801492 | 0.771565 | 0.794301 | 0.786169 | 0.787746 | 0.790577 | 0.79141 | 0.788357 | 0.792797 |
| Lung | 0.79608 | 0.77525 | 0.793123 | 0.794435 | 0.791987 | 0.78954 | 0.796107 | 0.790334 | 0.790282 |
| Muscle_Skeletal | 0.779067 | 0.759006 | 0.780461 | 0.775714 | 0.774085 | 0.775424 | 0.775399 | 0.770981 | 0.77295 |
| Esophagus_Mucosa | 0.792927 | 0.763298 | 0.785774 | 0.782096 | 0.779812 | 0.782891 | 0.787501 | 0.775147 | 0.777929 |
| Esophagus_Muscularis | 0.786048 | 0.766049 | 0.781259 | 0.783127 | 0.781263 | 0.779068 | 0.784338 | 0.777398 | 0.785203 |
| Whole_Blood | 0.808122 | 0.784989 | 0.803304 | 0.803906 | 0.800747 | 0.798743 | 0.804096 | 0.800218 | 0.80393 |
| Skin_Not_Sun_Exposed_Suprapubic | 0.782901 | 0.749986 | 0.773658 | 0.774382 | 0.774029 | 0.771202 | 0.778348 | 0.768952 | 0.773686 |
| Testis | 0.769392 | 0.7358 | 0.763324 | 0.759063 | 0.758056 | 0.756408 | 0.761967 | 0.761131 | 0.757602 |
| Adipose_Subcutaneous | 0.795779 | 0.772919 | 0.794064 | 0.790586 | 0.792146 | 0.789561 | 0.793822 | 0.787546 | 0.795928 |
| Cells_Cultured_fibroblasts | 0.795985 | 0.779762 | 0.794839 | 0.796589 | 0.794496 | 0.794541 | 0.796741 | 0.79003 | 0.792187 |
| Artery_Tibial | 0.790545 | 0.76756 | 0.785492 | 0.78004 | 0.778098 | 0.778498 | 0.786225 | 0.779509 | 0.781706 |
| Skin_Sun_Exposed_Lower_leg | **0.772106** | 0.740323 | 0.763365 | 0.761013 | 0.758437 | 0.760077 | 0.768233 | 0.759597 | 0.758128 |
| Thyroid | **0.776365** | 0.745303 | 0.769184 | 0.769032 | 0.768818 | 0.765115 | 0.762868 | 0.761224 | 0.765435 |
| Nerve_Tibial | **0.783039** | 0.759527 | 0.774918 | 0.774973 | 0.773309 | 0.772808 | 0.776014 | 0.771106 | 0.775442 |

## Methods

### Model architectures

The reference model architecture for the transformer is a stripped-down version of the Borzoi architecture (Linder et al. 2025). It comprises seven blocks of convolution and twofold max-pooling (yielding an embedding at 128bp resolution), four attention blocks, and then two rounds of UNet-style upsampling (yielding 32bp resolution), followed by separate output heads for each track.

We used a loss function consisting of a Poisson log-likelihood for the total counts for each output track, combined with a Multinomial log-likelihood for the count distribution over positions within each track, conditioned on the total counts for that track. With even weighting between these terms, this would constitute an independent Poisson loss at each position; however, we upweighted the multinomial term by a factor of four in an attempt to encourage correct total counts, for direct comparison to the loss used to train Borzoi (Linder et al. 2025).

Our convolutional-only model omits the attention blocks, while our Mamba and Hyena models replace the attention blocks with multiple layers of the appropriate architecture. The Hyena layers follow the architecture described in (Poli et al. 2023) and the Mamba layers as described in (Gu and Dao 2023) with the exception that, for both Hyena and Mamba, we developed bidirectional versions. At the time we developed these models, the published versions of Hyena and Mamba were “causal” architectures that only considered upstream flanking sequence as context, and not downstream sequence; this is appropriate for generative sequence modeling, where the next token to be emitted is probabilistically conditioned on previously-emitted tokens, but is inappropriate for seq2seq tasks where the output label at a given position is influenced by flanking sequence on both sides. To construct a bidirectional version of Hyena, it is sufficient simply to remove the filter padding from the Discrete-Time Fourier Transform so as to include flanking sequence on both sides. To make Mamba bidirectional, it is necessary to double up the model, processing the sequence in both forward and reverse directions in each layer. This approach was first described in Vision Mamba (Zhu et al. 2024); we added a reverse-complement operation for the backward scan and some parameter tying to approximate equivariance (specifically, the forward and backward projections into the expanded internal space of the SSM layer are tied). Subsequently to our development of these models, the developers of Hyena and Mamba have each released their own bidirectional modules within the PyTorch framework, the Mamba version being fully revcomp-equivariant (Hwang et al. 2024).

The Mamba architecture includes several hyperparameters: N is the dimension of the hidden state, and E is the expansion factor of the projection into the model’s internal space. In exploring the values of these and other architectural hyperparameters, we generally took the approach of increasing model size until available GPU memory was maxed out during training, since we observed GPU memory to be the limiting factor in model scaling. This coincidentally resulted in models with loosely comparable total parameter counts. For some models, notably (unstriped) Mamba and Hyena, accuracy began to drop off before this GPU memory limit was reached (due possibly to overfitting but perhaps also due to imperfect configuration of the optimizer’s learning schedule).

As well as the Mamba- and Hyena-based models, we also tried MultiResNet, a convolution-only architecture with shared parameters across scales, which is described in (Shi, Wang, and Fox 2023).

Table 1 lists all model architectures for which results are reported in this study; Figure 3 illustrates three of the best-performing configurations.

**Figure 3.**
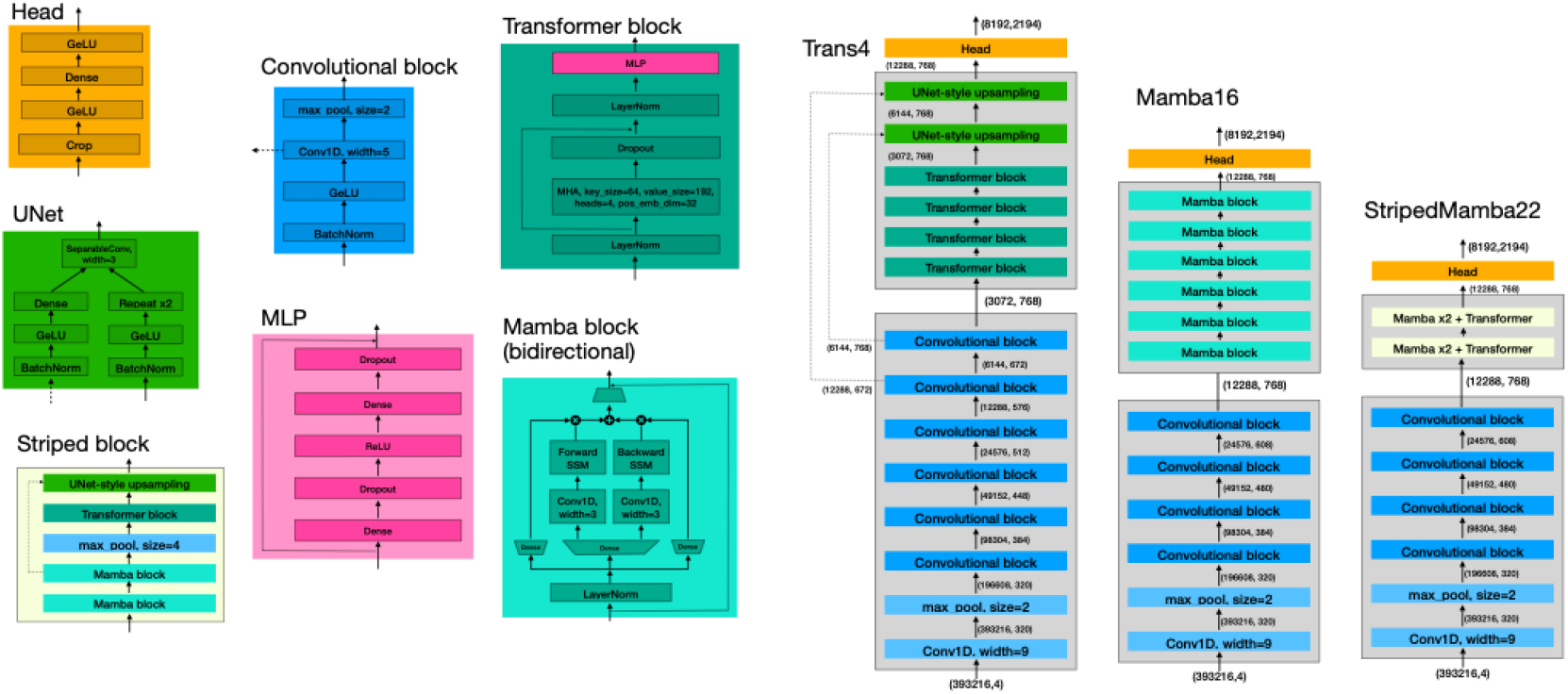
Architecture diagrams for three of the models evaluated in this study illustrate components used across different models.

### Model implementation

#### Framework

Our models and training algorithms are implemented in Jax, using the Flax library for convolution and attention.

#### Activation functions

We used SiLU activations for Hyena, and GeLU for all other models.

#### Normalization

We used BatchNorm with the convolutional layers, LayerNorm with attention and Hyena layers, and RMSNorm with Mamba layers. We observed that BatchNorm was unstable with Hyena (as has been reported with transformers).

#### Convolutional layers

All models began with at least five rounds of convolution; specifically normalization, activation, convolution, and twofold max-pooling, eventually expanding to a dimension-768 embedding over 32bp bins as shown in Figure 3. This convolutional stack converts an input tensor of size (393216,4) to an output of size (12288,768).

#### Transformer

The transformer architectures used 4 attention heads and a query/key dimension of 64. The transformer architecture introduced two additional convolution/pooling layers, bringing the bin size up to 128bp, followed by UNet-style upsampling for an eventual 32bp-resolution output. For the striped Mamba architectures, which involved fewer transformer layers and therefore required less stringent memory-reduction, the UNet wrapping each transformer involved just one 2x pooling and upsampling operation.

#### Flash attention

The Flash attention algorithm computes the attention matrix in chunks, so as to minimize the total transfers from the GPU’s high bandwidth memory to its shared memory cache (Dao et al. 2022). This requires low-level GPU access; for PyTorch this is implemented using CUDA. We employed a Flash-like attention-chunking schedule in Jax (P. Wang, n.d.) with the primary goal of reducing GPU out-of-memory errors by encouraging the Jax compiler to make fewer data transfers. This appears to be somewhat effective but does not guarantee optimal GPU memory performance; for such guarantees, a low-level GPU programming protocol such as Triton (Jax’s approach to low-level CUDA programming) would be required.

#### Positional encoding

We explored three strategies for encoding the distance between key and query locations in the attention computation: (1) the geometrically-spaced threshold indicator functions for relative feature separation as introduced by Enformer (Avsec et al. 2021), (2) Rotary Positional Embedding or RoPE (Su et al. 2021), (3) no positional encoding at all. For the Enformer-style encoding (1), we implemented a custom vector-Jacobian product (VJP), a.k.a. reverse-mode autodiff; however, we observed that this significantly slowed the implementation of Flash attention. We subsequently moved to RoPE (2), but (as noted in Results) found that RoPE provided no clear benefits when used with Mamba layers, so we then proceeded with no encoding (3).

#### Initialization

We used initializers as described in the Borzoi paper for attention and convolution layers (Avsec et al. 2021). For other layers, we generally employed Lecun initializers. When randomly initializing the weights of the SIREN function used for Hyena’s learnable convolution kernel, the typical approach is to maintain unit variance at the final layer (Sitzmann et al. 2020); however, when the generated function is used as a long convolution kernel for a length-L sequence, this leads to the convolution output having variance proportional to L, which causes exploding values when multiple Hyena layers are stacked at depth. Consequently, we found it useful to initialize the SIREN such that the final layer’s initial variance scales as 1/L.

#### Mamba associative scan

We implemented several versions of the Mamba associative scan algorithm, the best-performing of which was a recursive scan with gradient checkpointing.

#### Regularizers

We used L2 regularizers on all weights, with a loss multiplier of 10^−6^.

### Training data

All models were trained on 2,194 human genome-wide read-coverage datasets spanning 49 tissues and 4 types of experiment including 724 RNA-Seq and 321 DNase-Seq tracks from ENCODE (ENCODE Project Consortium 2012), 35 RNA-Seq tracks from GTEx via RECOUNT3 (Wilks et al. 2021), 434 3’ RNA-Seq tracks from the Tabula Sapiens project (Tabula Sapiens Consortium* et al. 2022), and 680 CAGE tracks from FANTOM5 (Noguchi et al. 2017). Of these, 1,694 tracks (including 680 CAGE, 580 RNA-Seq, and 434 3’ RNA-Seq tracks) were strand-oriented and partitionable into forward- and reverse-associated pairs of tracks.

The human genome was divided into 13,851 windows of size 393,216bp, with outputs pooled into 32bp; 2,048 32bp bins were cropped from each end prior to loss calculation. Coverage levels were squashed into 16-bit floating point values by the invertible monotonic transformation 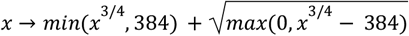. The set of 13,851 windows was partitioned into six folds, which were then grouped into a 4:1:1 train:validation:test split. We repeated all training experiments over four such data splits, with a different test set (and a different validation set) in each split. Training data were augmented by reverse complementation (with exchange of forward- and reverse-oriented tracks) and by shifting the sequence by up to 3 nucleotides in each window, replacing shunted-out bases with zeros in the one-hot encoded input tensors, while preserving the 32bp output bins unchanged.

### Training procedure

We used the Jax Optax framework’s implementation of the Adam optimizer, with the momentum decay rate parameters beta1=0.9, beta2=0.999. We used gradient clipping of 5.0 at the block (layer) level, and 10.0 at the global level. The learning rate was 0.0001 and we used 10,000 linear warm-up steps. In this paper, an “epoch” refers to a full iteration over the entire (shuffled) set of training windows, with a batch size of 2.

### Training hardware

Bilby models were trained on Google Cloud Platform g2-standard-96 nodes (96x 2.2GHz Intel Cascade Lake CPUs, 384Gb RAM, Debian Linux, 8x NVIDIA L4 GPUs each with 24Gb RAM, direct mounted disks).

Baskerville models were trained on comparable servers (64x 2.1Ghz Intel Xeon Gold CPUs, 250Gb RAM, Ubuntu Linux, 4x NVIDIA Titan RTX each with 24Gb RAM, networked filesystem). After confirming similar training convergence properties on both systems, the analogous total training time was estimated by comparison of epoch times over a small sample (~60 epochs). Each training run utilized only a single GPU.

### Benchmarking

#### Test set metrics

We first evaluated the models by averaging Pearson’s correlation coefficient (r) and the coefficient of determination (r^2^) across all output tracks and all windows in the withheld test set.

#### GTEx SNP classification

To assess the extent to which the models are identifying causal regulatory nucleotides, we further evaluated models by the GTEx classification benchmark described in (Linder et al. 2025). In this experiment, eQTLs from 49 tissues that had previously been fine-mapped using SuSIE and identified as causal with a posterior causal probability (PIP) of ≥0.9 were used as positives, with a negative set chosen to have marginal association significance with low PIP. In each case, the model is used to predict all output tracks for a window centered on the variant loci, for both reference and variant genotypes. The differences, and log-ratios, between reference and variant are computed. We then take the log-sum and L2-norm summaries of these differences and log-ratios, respectively, as described in (Linder et al. 2025). This is repeated for an ensemble consisting of models trained on all four data splits. The metrics we report for this SNP classification benchmark include area under ROC curve and Spearman’s rank correlation coefficient for the set of positives (i.e. the causal fine-mapped SNPs).

## Discussion

We found small but robust gains in prediction accuracy at a functional genomics task by using Mamba and Striped Mamba layers in combination with convolutional layers, compared to the state of the art convolution-attention architecture. Performance gains at the downstream SNP classification task are more mixed; the classifier performance of the Mamba-based methods is roughly equivalent to both the full Borzoi transformer model and the stripped-down version tested here (the best Mamba-based methods have, respectively, a mean AUROC within 0.007 of the transformer, a superior AUROC in 8 of 49 tissues, and a Spearman’s rank correlation coefficient that is greater by 0.006). For the configurations used here, which roughly match parameter counts and GPU memory usage, the Mamba-based models are also approximately comparable to the transformer models in training time.

Selective state space models like Mamba remain understudied compared to transformer models, and this potentially leaves considerable room for improvement on the results reported here. In our explorations of training hyperparameters, we lifted a number of elements wholesale from Borzoi, including the learning schedules, regularization schemes, and convolutional filters. Borzoi is a transformer-based model, and it is quite possible that some of these design choices are better suited to transformers than they are to Mamba. Perhaps even more pertinently, it bears noting that many of the models we evaluated here were unavailable in standard machine learning libraries, which necessitated that we develop our own versions (for which we chose to use Jax). It is entirely possible (and even likely) that the reference PyTorch methods include modeling optimizations that were unavailable to us at the time of writing. The landscape for such model development is changing rapidly; while we were writing this paper, the PyTorch ecosystem has advanced, with PyTorch-based libraries implementing the Borzoi models announced via preprint (Lal et al. 2024), while the developers of Mamba have released a bidirectional extension named Hydra (Hwang et al. 2024). Further, the period of this investigation saw the publication of several foundation models for genomic sequence based on Striped Hyena (Nguyen et al. 2023, 2024) and Mamba (Schiff et al. 2024; Fradkin et al. 2024), along with various Mamba-based models for proteins (Xu et al. 2024; Sgarbossa, Malbranke, and Bitbol 2024; Y. Wang et al. 2024).

With the caveat that the investigation described here must, consequently, be viewed as a preliminary one, we conclude that these results suggest a promising future for selective state space models as tools for supervised multi-task learning in functional genomics. For researchers interested in developing such models in the Jax framework, we further hope that our bilby source code repository may prove a useful resource.

## Acknowledgements

This project was supported by Calico Life Sciences LLC. IH received additional support from NIH grants R01GM080203 and R01HG004483.

